# Neural and autonomic predictors of future listening errors

**DOI:** 10.64898/2026.09.09.750180

**Authors:** Samuel S. Smith, Jenna A. Sugai, Kenneth E. Hancock, Daniel B. Polley

## Abstract

Some of the most demanding perceptual tasks require sustained vigilance to detect and classify infrequent, unpredictable targets within continuous sensory streams. Here, we designed a sustained listening task to ask whether cardinal classes of perceptual errors (misclassifications, miss, and false alarms) were associated with variability in event-related processing, ongoing cortical state, and autonomic arousal. We recorded 64-channel scalp EEG and pupil diameter in 39 adults while they monitored streams of random tone pips for occasional target sequences of repeating tones. A target-evoked negativity varied systematically with listening outcome: it was largest for correct reports, reduced for misclassifications, and nearly absent for misses. Notably, these differences emerged within the first 500 ms of the target sequence, before the information needed to classify its length was available. Related negativities also accompanied false alarms despite the absence of a target. By contrast, neural synchronization to acoustic cues in individual tone pips did not differ across perceptual outcomes. We then looked further back in time, to the 1 s period preceding target onset, and identified additional signatures of forthcoming misclassifications and false alarms in pupil diameter and EEG aperiodic slopes. Models evaluated with leave-one-subject-out cross-validation showed that baseline state measures and event-related negativity amplitude provided complementary information about the probability and type of forthcoming errors. These findings link perceptual errors during sustained listening to higher-order cortical and arousal-state dynamics rather than fluctuant encoding of bottom-up acoustic cues.

## INTRODUCTION

Perceptual decision making is a dynamic process that relies on monitoring, identifying, and responding to statistical patterns conveyed via our sensory systems. This process is prone to errors. Misinterpretations, misses, and false reporting of stimuli are all common outcomes that can alternate from one moment to the next. While sensory processing is traditionally framed in terms of stimulus-response relationships, the mapping from a single sensory input to disparate perceptual failures highlights the influence of top-down extra-sensory factors in shaping our awareness of the sensory environment.

Top-down contributions are evident in how the brain tracks sensory regularities. For example, scalp-EEG negativities associated with auditory structure tracking (Barascud et al. 2016; Sohoglu and Chait 2016; Southwell et al. 2017) are modulated by diverse non-sensory factors including auxiliary task performance (Herrmann and Johnsrude 2018), feature-directed attention (Ringer et al. 2023), task familiarity (Herrmann et al. 2021), short-term memory load (Hu and Chait 2026), healthy aging (Herrmann et al. 2019) and neuromodulation (Al Jaja et al. 2020). Crucially, while such event-related activity is typically considered in error-free contexts, its sensitivity to higher-order dynamics suggests it may reflect perceptual instability during listening.

Brain dynamics occurring prior to stimulus onset can also modulate cortical processing of forthcoming sensory stimuli (McCormick et al. 2020). Baseline state variables, such as aperiodic neural activity, quantify cortical synchrony and excitation, and capture perceptual variability surplus to sensory input (Eldar et al. 2013; Waschke et al. 2019; Donoghue et al. 2020; Waschke et al. 2021; Parajuli et al. 2023). In addition, fluctuations in global arousal, commonly captured via pupil tracking, tightly couple suboptimal behavioral performance at both low and high extremes across multiple task domains (McGinley et al. 2015; Beerendonk et al. 2024; de Gee et al. 2024; Hulsey et al. 2024; Smith et al. 2025).

Together, these extra-sensory processes occurring before and during sensory encoding demonstrate that perceptual fluctuations are not the product of stochastic internal noise but instead arise from structured and empirical brain dynamics. Yet, whether these neural systems act in concert or operate as independent pathways remains unclear. Furthermore, behavioral errors are often treated as a single phenomenon, with cued, forced-choice paradigms grouping distinct failure modes under a single incorrect label. Yet, misinterpreting sensory information is fundamentally distinct from failing to detect it, and both differ from reporting a stimulus in its absence.

Here, we developed a continuous auditory monitoring task that captures this diversity of human errors, incorporating both detection and discrimination of uncued targets during two-minute bouts of active listening without trial segmentation. By simultaneously tracking scalp-EEG and pupillometry, we measured the respective contributions of evoked activity and baseline brain states to perceptual errors. We hypothesized that distinct, canonical listening errors (misclassification, misses, and false alarms) reflect the breakdown of dissociable stimulus-evoked and state-dependent pathways. Finally, we built a cross-validated decoding framework, demonstrating that these signatures of monitoring failure are generalizable across individuals.

## MATERIALS AND METHODS

### Participants

All procedures were approved by the Mass General Brigham Institutional Review Board (#IRB2019P002423) and took place at Mass Eye and Ear. Participants were recruited through flyers, word of mouth, and by posting to the Mass General Brigham participant recruitment website. After providing informed consent, study participants between the ages of 18 and 50 were screened for English fluency, cognitive function, medical history and mental health. Hearing status was also assessed via otoscopy and through pure tone audiometry by a licensed audiologist. Participants were excluded if: 1) the air-conduction threshold exceeded 25 dB HL at any frequency from 0.25 to 2 kHz in either ear; 2) the interaural threshold difference exceeded 10 dB at each of three contiguous frequencies. Because the task required continuous pupillometry across 2-minute runs, participants also underwent screening for blink frequency and pupil data quality following an initial familiarization block. In total, 41 study participants passed all screening criteria and completed the full testing blocks. Of these, 2 participants were excluded due to outlier performance on the task, with average correct target classification below 50% accuracy. The remaining 39 participants comprised the primary analysis cohort, with further restrictions to sample sizes due to excessive measurement noise or error type absences detailed below.

### Task

Participants completed 20 runs of a listening task that required continuous monitoring of two-minute-long random tone clouds. Tone clouds consisted of tones with carrier frequencies drawn from a set of 25 equally spaced frequencies in the log domain, covering a two-octave span from 353.6-1414.4 Hz. Tones were amplitude modulated at 40 Hz and presented every 125 ms (8 Hz) with a duration of 100 ms each. Tones were presented diotically through insert ear headphones (EarTone 3A, Oaktree Products, Chesterfield, MO) at an intensity of 75 dB SPL. Intermittently, unprompted targets in the form of patterns of consecutive repeating tones were embedded within the tone cloud.

The target patterns consisted of either 4, 5, or 8 tones in a row (probabilities of 0.5, 0.25, and 0.25, respectively) at carrier frequencies of either 500 Hz or 1000 Hz. Targets at each carrier frequency were presented at random intervals, each initially drawn from a uniform 0 to 10 second distribution. A 6-second deadtime was enforced upon each target onset, and target events could never have the same carrier frequencies more than twice in a row, resulting in the time between any two targets ranging from 6 to 20 seconds, with an approximately triangular distribution. Upon detecting a target, participants were asked to report its length as either 4 or more than 4 tones-in-a-row via button press. The outcomes of button presses were either correct classification, misclassification, or false alarm (i.e., when no target was presented). A target could not be accurately classified as four vs. more than four within the initial 500 ms (i.e., until the first four tones had passed). A response window was open until 2.5 seconds post-target onset, with feedback provided 0.5 seconds after each button press, lasting for 1.5 seconds. If no button was pressed in the response window, this was classed as a miss.

To ensure task competency, participants first completed runs with only targets (no tone cloud), followed by a familiarization block with four runs of the full task (B0), followed by four test blocks of four runs each (B1-B4; **Fig. 1A**). All testing was performed in a sound attenuating RF-shielded booth measuring 2.9 m x 2 m x 2 m (L x W x H; Eckel Industries Inc., Ayer, MA) with an ambient noise level less than 16.4 dBA. Participants sat in low-light isoluminous conditions and fixated towards the center of the front-facing monitor while minimizing blinking.

**Figure 1.**
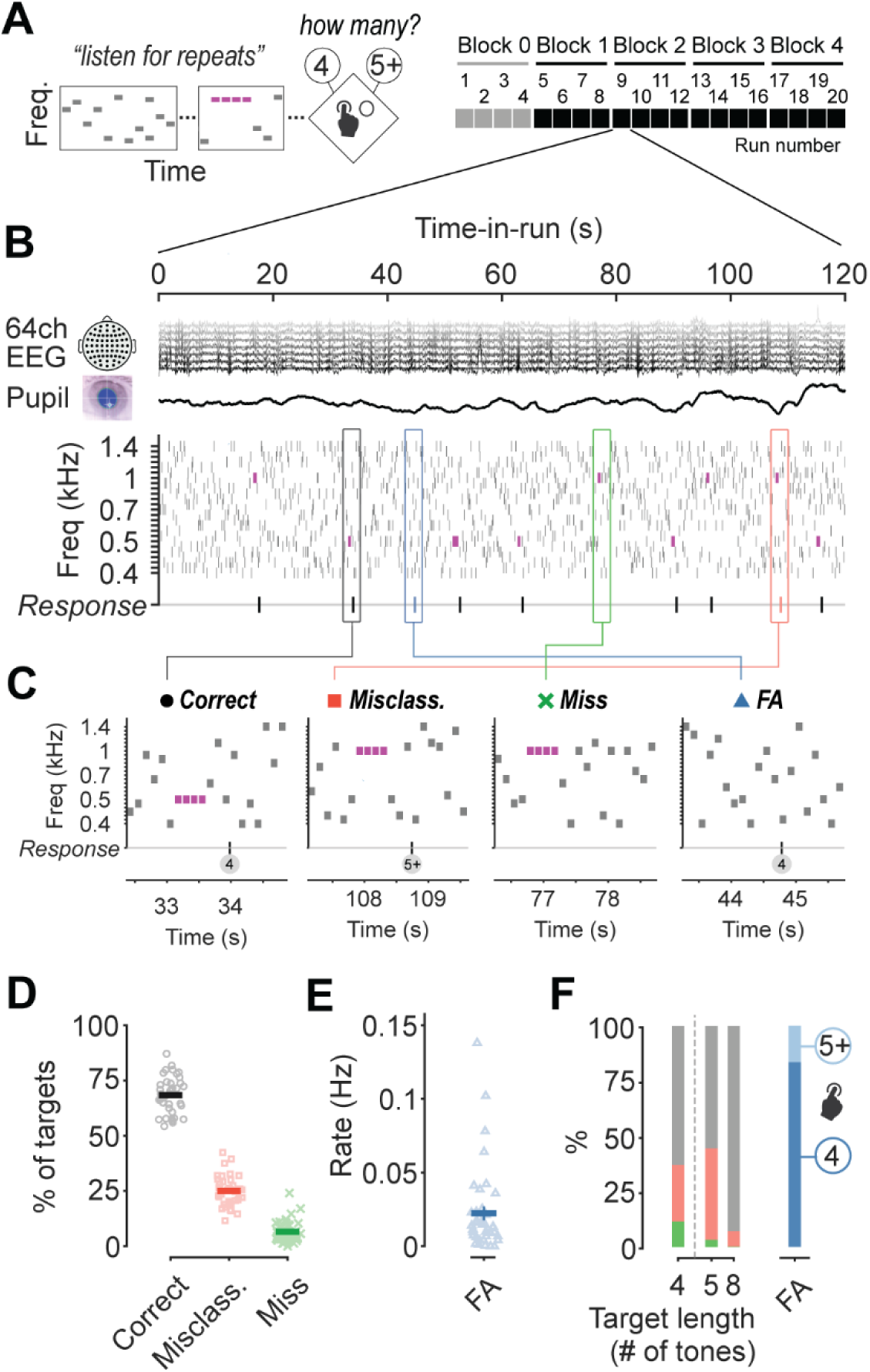
Multiple error types occur during sustained monitoring. **A**, Participants monitored random tone sequences and reported targets by identifying repeating patterns. Upon detecting a target (≥4 consecutive identical tones), they classified repeat counts via a keypad. The study consisted of 5 blocks of 4 runs each, with the initial block (B0) providing task familiarization. **B**, Neural and autonomic dynamics were monitored during task performance with 64-channel scalp EEG and infrared pupillometry. An example 2-minute run illustrates the continuous stream of tones with the occasional appearance of repeating targets (purple) and behavioral responses. **C**, Aside from correctly reporting a target (far left), participants could make three distinct error types: misclassifications (center left, incorrectly reporting the number of repeating tones), misses (center right, failing to report a target altogether), and false alarms (far right, mistakenly identifying a non-existent pattern as a target). **D**, The likelihoods of correctly reporting a target, misclassification, and misses are shown for each participant during test blocks. Group means ± SEM are overlaid. **E**, False alarm rates are presented for study participants. Group mean ± SEM is overlaid. **F**, Stacked bar plots illustrate the distribution of behavioral outcomes across target repeat lengths (left) and the mean proportion of false alarm presses for each available button (right).

### EEG preprocessing

EEG data was recorded using a 64-channel electrode montage at a sampling rate of 8192 Hz with electrodes on the left and right mastoids and on the lateral side and underneath each eye (BioSemi ActiveTwo system, Wilmington, NC). Preprocessing was performed using EEGLAB (Delorme and Makeig 2004), NoiseTools (de Cheveigné and Simon 2008), and custom scripts in MATLAB (MathWorks, Natick, MA). Offline, raw data was downsampled from 8192 Hz to 256 Hz. Bad channels were detected via the clean_rawdata function on traces highpass filtered at 1 Hz (this filtering step was applied exclusively for bad channel detection). The clean_rawdata function was set to detect flatline channels, uncorrelated channels, and line noise (using default parameters). Next, a conservative 0.3 Hz high-pass filter was applied using a zero-phase FIR design to eliminate low-frequency drift while preserving temporal precision, followed by removal and spherical interpolation across the previously determined bad channels. Blinks were removed from the EEG data using denoising source separation (3 components; de Cheveigné and Simon, 2008). Channels were re-referenced against mastoid recordings. The data were then epoched ±2.5 seconds around target onsets, false alarms, and randomly sampled correct rejects (with a total of 32 non-target trials per run). Trials with more than 10 channels exhibiting values greater than 120 μV were removed. If fewer than 10 but at least one channel exceeded this threshold, these channel(s) were replaced with projections based on neighbors using spherical splines.

Additionally, trials that deviated from the participant mean by more than 5 standard deviations were removed, resulting in an average of 2.7% of trials removed. One participant’s EEG data was excluded due to excessive missing values (>20%). For ERP analyses only, a 125 ms moving mean was applied to attenuate 8 Hz tone-rate modulations.

### Auditory steady-state response (ASSR) analysis

ASSR spectral power at the 40 Hz amplitude modulation rate was computed within both the 0.5 s evoked window and the 1 s baseline window. For the baseline window, estimates were computed using a 0.5 s sliding window across the full 1s period and averaged, to match the spectral resolution and SNR characteristics of the evoked-window measure. For both windows, trials were averaged in the time domain prior to Fourier analysis to isolate stimulus-locked activity at 40 Hz while minimizing non-stimulus-related variation. We implemented a bootstrapping procedure to equate trial counts across error-outcome contrasts. For each contrast, trials were randomly subsampled to the minimum available trial count across conditions, with this process repeated 100 times and the resulting Fast Fourier Transform measures averaged. To quantify signal-to-noise ratio (SNR), we calculated log-power differences between 40 Hz and the adjacent off-frequencies.

### Aperiodic slope analysis

We characterized the aperiodic component of the EEG power spectrum on a single-trial basis in 1-second baseline windows. For each trial, power estimates were computed via Welch’s method (Welch 1967). We then fit a first-order polynomial to the log-transformed power in the 2–50 Hz frequency range using ordinary least squares regression. To prevent prominent alpha band oscillations from biasing the aperiodic fit, 8–12 Hz was excluded from the fitting procedure. The slope of this fit constituted each trial’s aperiodic slope value, which was carried forward for subsequent statistical analysis.

### Pupillometry

Pupil dynamics were recorded using an EyeLink 1000 system (SR Research) sampled at 1000 Hz. Eye blinks were automatically detected by the system and corrected using linear interpolation. The resulting pupil traces were smoothed using a forward-only, low-pass Butterworth filter with a 0.6 Hz cutoff frequency. Baseline pupil diameter was defined as the mean value calculated within a 1 s window immediately prior to target onset. Trials were excluded from subsequent analyses if more than 75% of the data points within this baseline window were missing due to blinks or artifacts, resulting in an average rejection rate of 18% of trials. To examine nonlinear relationships, baseline pupil diameters were discretized into five proportionally split rank-ordered bins (quintile 1 through 5) following established protocols (Beerendonk et al. 2024; Smith et al. 2025). We derived U-indices by subtracting error likelihood during intermediate pupil-linked arousal (Q3) from the mean error likelihood during the lower and upper extremes (Q1 and Q5).

### Statistical analyses

All statistical tests evaluated significance at α=0.05. Behavioral error rates were computed across the full sample (N=39). For analyses contingent on specific error types, sample sizes were restricted to participants with available error trials (N=38 for misses; N=37 for false alarms). As detailed above, one participant was excluded from EEG analyses due to excessive noise artifacts.

To assess the influence of target length on behavioral performance, error likelihoods were analyzed with one-way repeated-measures ANOVAs followed by post-hoc pairwise comparisons with Holm-Bonferroni corrections. One-sample *t*-tests were used to evaluate differences in the proportion of respective button presses (reporting 4 tones vs. 5+ tones) during false alarms.

Because EEG analyses involved comparisons across 64 channels, and in some analyses across multiple time points, family-wise error rate was controlled using a non-parametric cluster-based permutation procedure implemented in the FieldTrip toolbox that were applied to the ERP, ASSR, and aperiodic slope analyses (Maris and Oostenveld 2007; Oostenveld et al. 2011),. Default parameters were applied using the max-sum *t*-statistic criterion, with clusters evaluated against a two-tailed significance threshold (α = 0.05 critical threshold, *p* < 0.025 per tail). For ERP analyses, cluster permutations were evaluated over a 0–0.5 s evoked window aligned to target onsets, and separately for a –1–0 s baseline window. For false alarms, baseline and evoked analysis windows were aligned relative to pseudo target onsets estimated by subtracting each participant’s mean response time from their button press timestamps. For analysis of 40 Hz envelope following response SNRs and aperiodic slopes, cluster permutations were evaluated across channels only. All analyses were performed at the group level, across participants using individual average contrast maps, with participant identities randomly shuffled across iterations (1000 iterations). For each of these three analyses, pairwise contrasts among the three target-present outcomes (misclassification vs. correct, miss vs. correct, and misclassification vs. miss) were additionally Holm-Bonferroni-corrected as a family. Because false alarms arise from separate target-absent trials, this contrast (false alarm vs. correct rejection) was evaluated separately.

For tonic pupil U-indices, the correct-trial U-index is fully determined by the misclassification and miss indices (U_correct_ = −[U_misclassification_ + U_miss_]). We therefore tested only the misclassification and miss U-indices against zero, using one-sample *t*-tests Holm-Bonferroni corrected as a family of two. As above, the false alarm U-index was evaluated in a separate one-sample *t*-test.

To evaluate the predictive capacity and functional independence of baseline and evoked measures, we employed out-of-sample trial-level forecasting using leave-one-subject-out cross-validation (LOSO-CV). Multivariate logistic regression models were trained on N-1 participants to predict upcoming trial-level error outcomes (misclassifications, misses, and false alarms) on held-out test participants. Features were z-scored within participants. Baseline aperiodic slopes and ERP amplitudes were first averaged across previously identified spatiotemporal clusters, and baseline tonic pupil size was squared prior to normalization. Model performance was evaluated as the difference in true error rates between trials assigned to high- and low-risk probability pools based on within-participant upper and lower quintile cutoffs (top/bottom 20th percentiles). As a robustness check, this percentile threshold was systematically varied from 1% to 99% to assess how prediction margins scaled with the stringency of the decision criterion (**Fig. S4**). For each error type, performance was compared across four conditions: the intact joint model, baseline-shuffled, evoked-shuffled, and a fully shuffled empirical null (100 iterations per subject), using a one-way repeated-measures ANOVA. Three contrasts, Holm-Bonferroni-corrected within each error type and reported via one-sample *t*-tests, assessed whether the intact model exceeded the null and whether shuffling either feature domain significantly degraded performance relative to the intact model. To rule out shared dependencies with time, we repeated this procedure using an expanded model that additionally incorporated session time, run time, and inter-target interval as features. For each error type, we compared the expanded model’s performance to a version in which both baseline and evoked features were shuffled. We ran one-sample t-tests to determine whether baseline and evoked features had predictive value beyond these time-related factors.

## RESULTS

### Multiple error types occur during sustained monitoring

In this study, 39 participants were presented with a continuous stream of tone pips while neural and autonomic activity was simultaneously recorded with 64-channel scalp EEG and pupillometry.

Participants completed five blocks in total, consisting of one familiarization block and four test blocks, with each block comprised of four two-minute runs (**Fig. 1A**). The experiment was designed as a sustained vigilance task that challenged participants to detect occasional targets in the form of low (0.5 kHz) or high (1 kHz) frequency tones repeated at least four times (**Fig. 1B**). Upon detecting a target, participants reported the number of repeat tones via a keypad (either 4 or more than 4). To encourage vigilance and minimize cuing effects, targets occurred infrequently (∼0.08 Hz) with unpredictable timing.

This design created a challenging listening task (mean accuracy=68.4%, SD=8.2%; **Fig. 1C-D**) characterized by three classes of behavioral errors. In the presence of a target, errors occurred either as misclassifications, where participants detected a target but incorrectly reported the string length (mean=25.0%, SD=6.7%), or as misses, defined by the absence of reporting during the response window (closing 2.5 seconds post-target onset; mean=6.6%, SD=4.9%). Conversely, in the absence of a target sequence, errors occurred as false alarms, defined as an erroneous keypad press (mean rate=0.022 Hz, SD=0.029 Hz; **Fig. 1E**).

Beyond their behavioral definitions, these error classes reflect different processing failures. Misclassifications capture successful target detection but a failure in length discrimination. Consistent with a boundary-decision error, misclassifications occurred most frequently for stimulus lengths closest to the classification boundary (i.e., lengths of 4 or 5 tones; one-way RM-ANOVA, *F*_(2, 76)_ = 100.6, *p* < 0.001; post hoc comparisons, *p* < 0.001 for all contrasts; **Fig. 1F**). In contrast, misses represent a failure in detection, occurring overwhelmingly during the shortest target strings (*F*_(2, 74)_ = 57.5, *p* < 0.001; post-hoc comparisons, *p* < 0.001 for all contrasts). False alarms instead represent errors of commission, with participants more likely to falsely report shorter rather than longer sequences when no target was present (paired t-test, *t*_(36)_ = 16.94, *p* < 0.001).

### Magnitude of target-aligned ERP scales with type of performance failure

To investigate potential neural drivers of variability in performance errors, independent of stimulus identity, we examined scalp EEG activity during a time-locked window spanning the first four tone pips (500 ms) of target sequences. As all targets contained at least four pips, this window provided a strictly controlled comparison point before distinguishing information regarding target length was available.

A prominent ERP negativity emerged across the scalp when the EEG was aligned to target onset, with cluster-based permutation testing showing a significant difference from baseline (*p*_cluster_ < 0.001), with the underlying negativity maximal over central electrodes (**Fig. 2A**). Partitioning trials by subsequent behavioral outcome revealed that the magnitude of this negativity scaled with error outcome. The ERP was attenuated for misclassifications relative to correct trials (*p*_cluster_ = 0.016; **Fig. 2B-C**, **Fig. S1**) and nearly abolished for misses relative to correct trials (*p*_cluster_ = 0.021). This reduction was also significantly more pronounced for misses than for misclassification (*p*_cluster_ = 0.012). As by definition false alarms arise in the absence of a target, we assessed this error separately. A negativity was observed on false alarm trials relative to randomly sampled correct rejects (*p*_cluster_ = 0.002; **Fig. 2E-F**), suggesting that this ERP provides a general signature of a forthcoming perceptual error, regardless of whether a target was physically presented.

**Figure 2.**
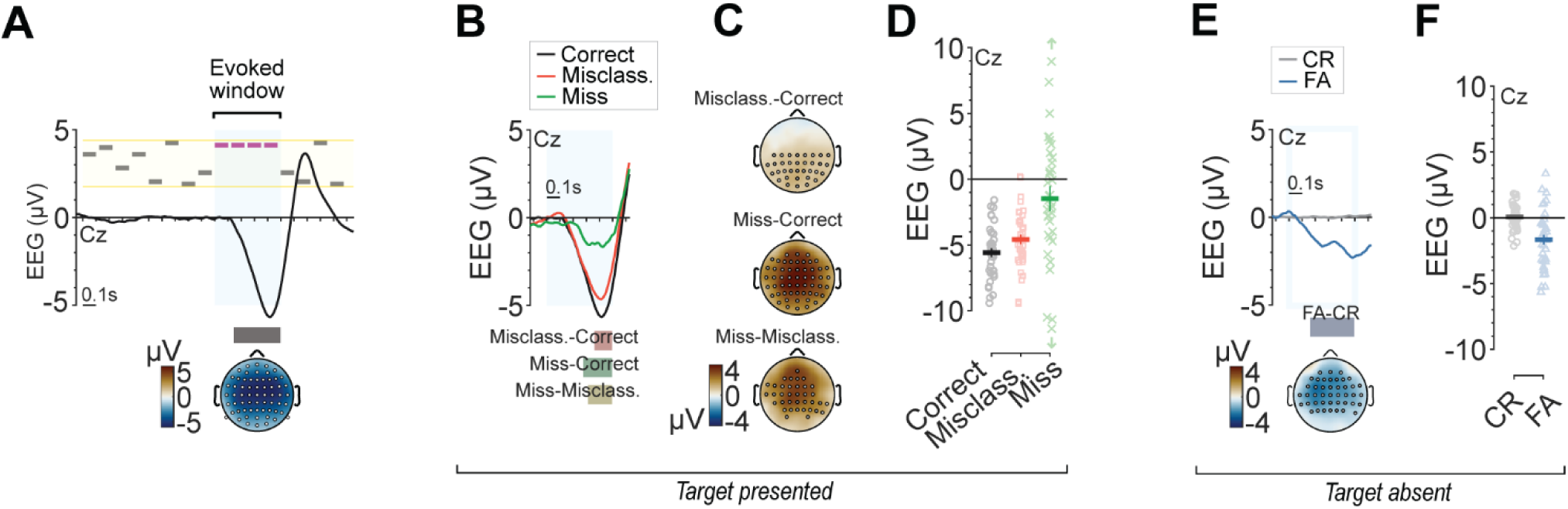
Magnitude of target-aligned ERP scales with type of performance failure. **A**, Schematic of the analysis window during target monitoring, with the grand-average event-related potential (ERP) at electrode Cz (black trace) overlaid. The thick black bar beneath the trace and the white dotted markers on the scalp topography indicate, respectively, the temporal extent and sensor distribution of the cluster identified as part of the cluster-based permutation analysis. **B**, Grand-average ERPs at Cz for target presentations, stratified by behavioral outcome: correct reports (black), misclassifications (red), and misses (green). Horizontal bars indicate temporal windows of identified clusters for each of the three pairwise contrasts. **C**, Scalp topographies of differential ERP amplitude for each of the three pairwise error contrasts, with dotted markers indicating the sensor distribution of identified clusters. See **Fig. S1** for full spatiotemporal profiles. **D**, Individual-subject ERP amplitudes at Cz for target presentations, stratified by behavioral outcome (colors as in B). Group means ± SEM are overlaid. Upward/downward arrows indicate values outside the plotted range. **E**, Grand-average ERPs at Cz in the absence of targets, stratified by behavioral outcome: correct rejections (grey) and false alarms (blue). The horizontal bar indicates the window of the identified cluster-level difference between conditions. The accompanying scalp topography shows the corresponding differential ERP amplitude, with dotted markers indicating the sensor distribution of the identified cluster. **F**, Individual-subject ERP amplitudes at Cz in the absence of targets, stratified by behavioral outcome (colors as in **E**). Group means ± SEM are overlaid.

We evaluated whether this signature might be confounded by task-related factors. We performed several alternative contrasts and sanity checks, including carrier tone frequency, target length, session time, run time, and inter-target intervals. Cluster-based permutation testing revealed no significant differences across any of these secondary factors (*p*_cluster_ > 0.05 for all; **Fig. S2**).

### Early auditory steady-state responses remain stable

To determine whether these ERP differences reflect higher-order cognitive processing rather than lower-level deficits in sensory encoding, we evaluated auditory steady-state responses (ASSR). Because all acoustic stimuli were amplitude modulated at 40 Hz, ASSR spectral power during the same 500 ms time-locked window provided a direct index of stimulus encoding quality.

Robust 40 Hz envelope-following responses were evident across the scalp (*p*_cluster_ < 0.001, **Fig. 3A-B**). However, once trial counts were matched across behavioral outcomes, 40 Hz signal-to-noise ratio (SNR) did not significantly differ between any pairwise contrasts comparing between outcomes (*p*_cluster_ > 0.05 for all, **Fig. 3C-D**). The stability in sensory encoding suggests that the modulation of target-aligned ERPs by trial outcome reflects top-down influences rather than degraded neural synchronization to the bottom-up acoustic cues.

**Figure 3.**
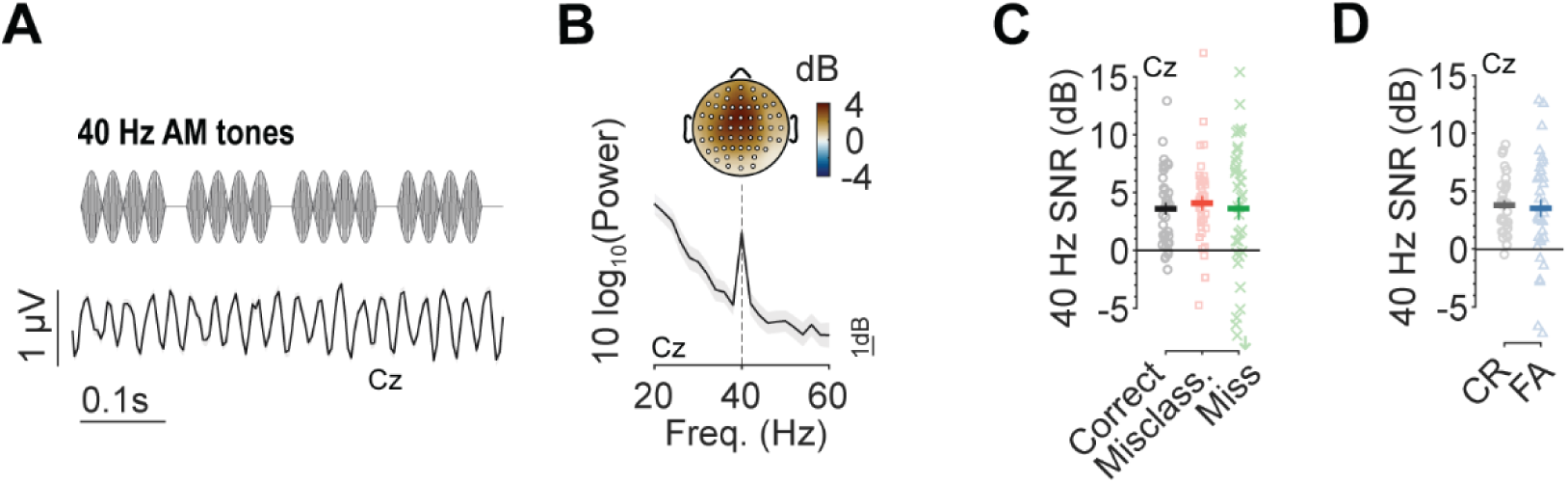
Early auditory steady-state responses remain stable. **A**, Grand-average time-domain envelope-following response at electrode Cz, elicited by the 40 Hz amplitude-modulated tone pips. For visualization only, the signal was high-pass filtered (30 Hz cut-off) to isolate the response. **B**, Power spectrum at electrode Cz showing a sharp peak at the 40 Hz modulation frequency. Scalp topography shows the corresponding 40 Hz signal-to-noise ratio (SNR), which was maximal over centro-frontal sensors. **C**, Individual-subject 40 Hz SNR values at electrode Cz for the three target-presented outcomes: correct reports (black), misclassifications (red), and misses (green). Group means ± SEM are overlaid. Downward arrow indicates value below the plotted range. **D**, As in **C**, for trials without a target present: correct rejections (grey) and false alarms (blue).

### Baseline cortical states are linked to specific performance failures

Having established that neural activity evoked by the initial buildup of target awareness – but prior to the availability of target length information – is linked to performance errors, we next examined the period preceding the onset of the target itself. Endogenous fluctuations in baseline cortical states continuously shape sensory-guided behavior. Do processing differences originating from baseline cortical states also shape upcoming errors?

To address this, we employed two assays of cortical state within a one-second window directly preceding target onset (**Fig. 4A**). As an index of cortical state, we measured the aperiodic slope of the single-trial EEG power spectrum. Steeper aperiodic slopes have been interpreted in prior work as reflecting a shift toward stronger cortical inhibition and increased neural synchronization, features previously linked to tradeoffs between sensory detection and discrimination performance (Donoghue et al. 2020; Waschke et al. 2021; Parajuli et al. 2023). Pairwise cluster-based permutation tests among the three target-present outcomes showed that misclassified targets were preceded by significantly steeper aperiodic slopes than correct trials (*p*_cluster_ = 0.021), that were prominent over parieto-central scalp regions (**Fig. 4B-D**). Neither the miss vs. correct nor the miss vs. misclassification contrasts yielded significant differences (i.e., *p*_cluster_ > 0.05 for both). False alarms were also preceded by significantly steeper aperiodic slopes relative to correct rejections (*p*_cluster_ = 0.015), that were prominent over more frontal regions (**Fig. 4E-G**).

**Figure 4.**
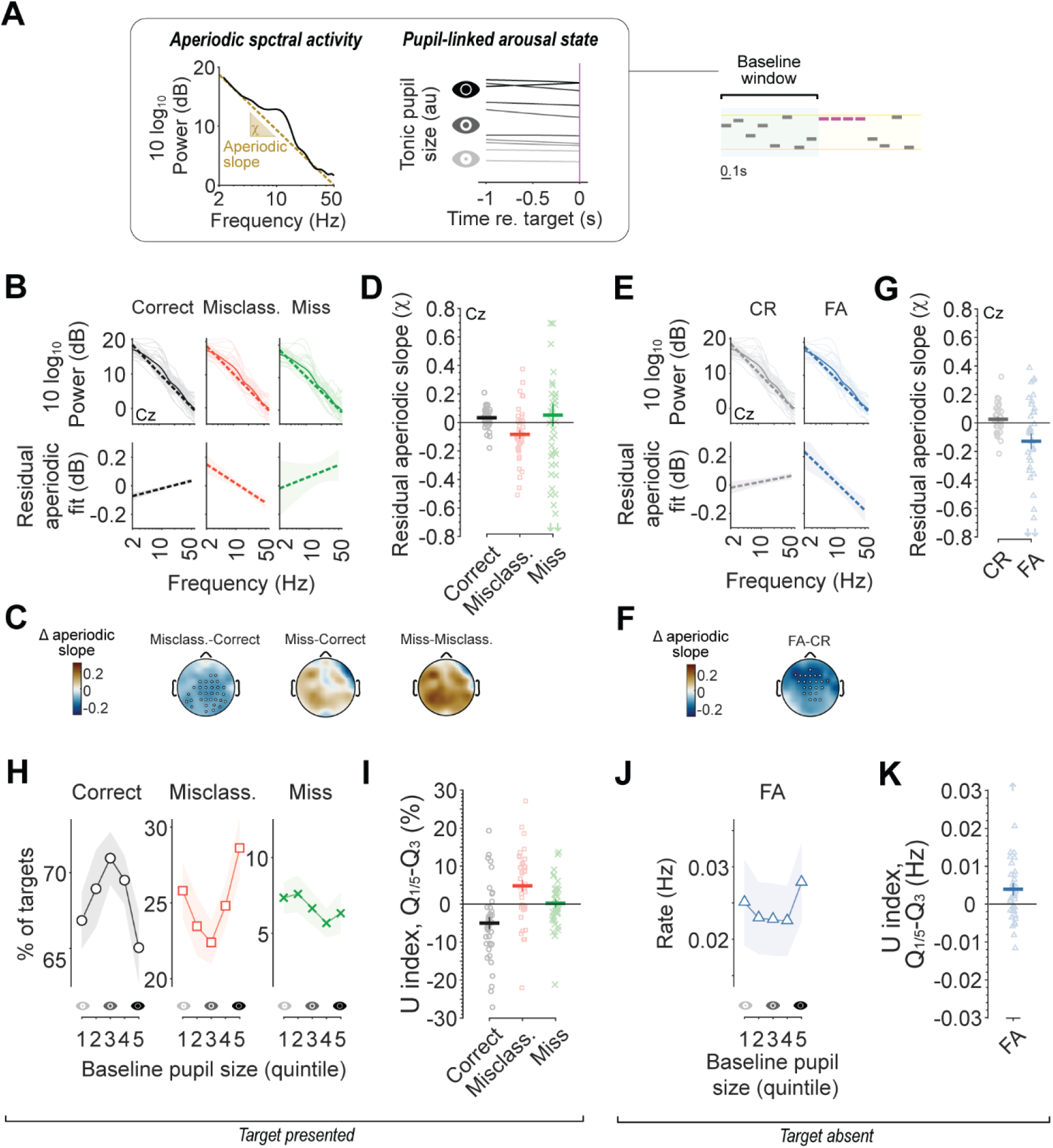
Baseline cortical states are linked to specific performance failures. **A**, Schematic illustrating the 1 s baseline window, during which we measured two proxies of cortical state: the aperiodic slope of spectral activity and pupil-linked arousal. **B**, Top: baseline power spectra at electrode Cz, stratified by behavioral outcome and fitted with aperiodic slopes. Dark and light lines represent the sample mean and individual participants, respectively. Bottom: the same aperiodic fits after subtracting each participant’s own mean slope across conditions, isolating within-subject differences between outcome types. **C**, Scalp topographies showing differences in aperiodic slope between error contrasts, with sensor clusters indicated by dots. **D,** Individual-subject residual aperiodic slopes at electrode Cz for the three primary outcomes when a target was presented: correct reports, misclassifications, and misses. Group means ± SEM are overlaid. **E-G**, as **B**-**D** for target-absent outcomes of CR and FA. Upward/downward arrows indicate values outside the plotted range. **H**, Relationship between baseline pupil size (binned into within-participant quintiles) and error likelihood, for the three target-present outcomes: correct reports (black), misclassifications (red), and misses (green). Shading indicates ± SEM. **I**, U-indices capturing the change in error likelihood at the lowest and highest extremes of pupil-linked arousal relative to intermediate arousal (mean[Q1,Q5] − Q3), computed for correct trials, misclassifications and misses. Note, the U-index of correct trials is mathematically fully determined by the misclassification and miss indices. **J-K**, As in **H**-**I**, for false alarms during target-absent periods.

We also assessed the influence of global arousal state within the baseline window, indexed by pupil size in relation to performance. We observed non-monotonic effects, with correct trials demonstrating a classic inverted-U relationship (**Fig. 4H–I**). Underlying this, pupil-linked arousal showed a significant quadratic relationship with misclassifications (one-sample *t*-test on U-indices against zero: *t*_38_ = 3.21, *p* = 0.005), but not misses (*t*_37_ = 0.19, *p* = 0.85). In the absence of a target, false alarms also showed a significant quadratic relationship with baseline arousal (*t*_36_ = 2.39, *p* = 0.022; **Fig. 4J-K**). Misclassification and false alarm errors were more likely during either hypo- or hyper-aroused states, whereas optimal performance occurred during moderate baseline arousal.

Together, these two markers of baseline cortical state were jointly associated with subsequent error outcomes. Given this, we evaluated whether these effects might be related to differences in time-locked scalp EEG activity within the same baseline window. However, we observed no such differences, nor in 40 Hz envelope-following responses, across error types during the 1 s period prior to target onset (*p*_cluster_ > 0.05 for all, **Fig. S3**).

### Trial-level prediction decouples evoked dynamics from baseline states

While baseline states and evoked responses were both associated with behavioral errors, the evoked ERP dynamics could be a product of the preceding baseline brain state dynamics, rather than an independent contributor. To address this, and to test whether these measures predict forthcoming errors, we trained logistic regression models to predict error outcomes in held-out participants (**Fig. 5A**). To capture each model’s predictive utility, we ranked held-out events by estimated error probability and compared true error rates between the upper and lower 20^th^ percentile (i.e., high and low risk) pools. For each error type, this measure was then assessed across the intact joint model, baseline-shuffled, evoked-shuffled, and a fully shuffled null model.

**Figure 5.**
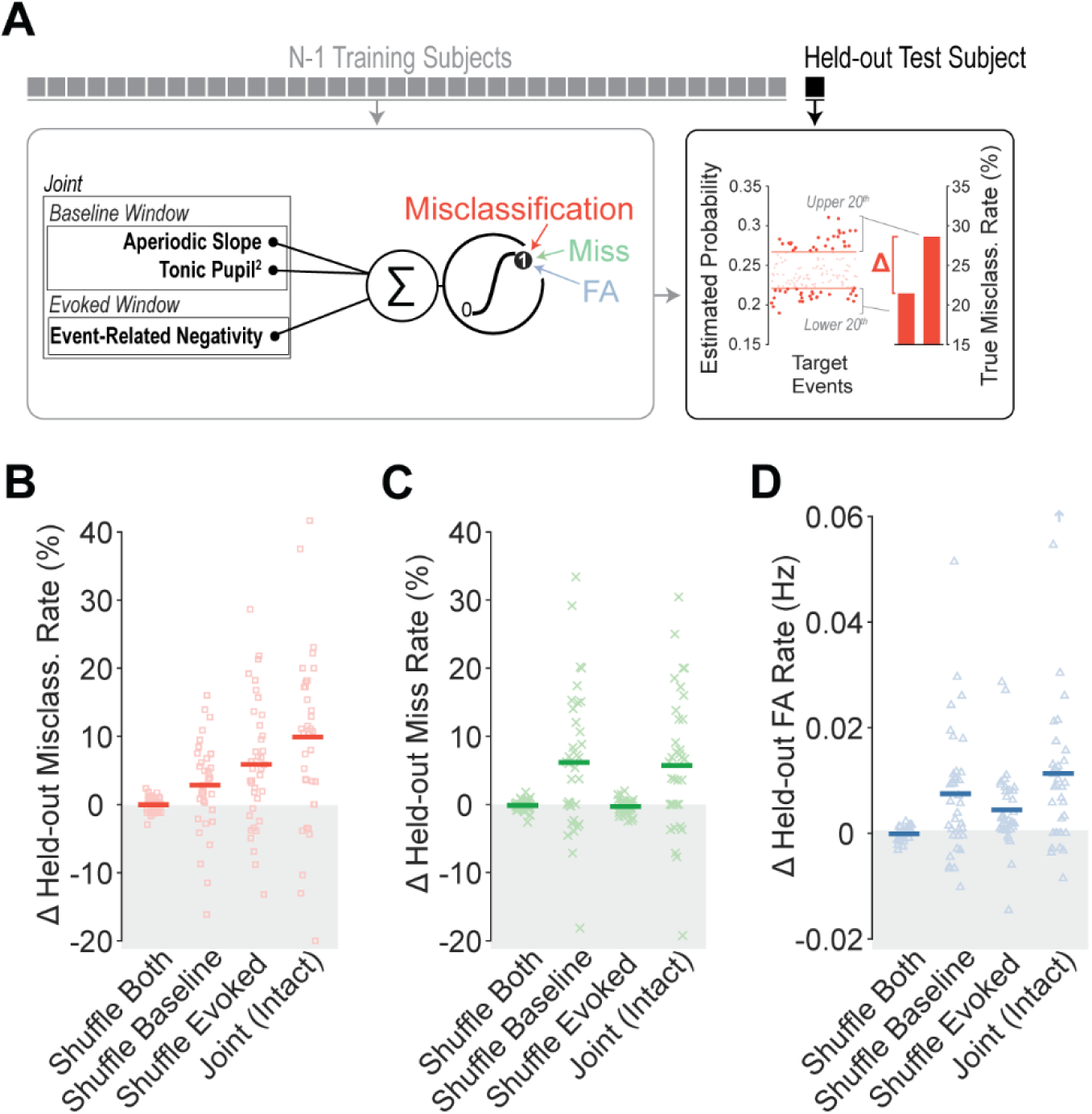
Trial-level prediction decouples evoked dynamics from baseline states. **A**, Multivariate logistic regression models were trained in a leave-one-subject-out design to forecast trial-level error likelihoods. To assess practical predictive utility, trials were assigned to “likely” or “unlikely” error pools based on forecast probabilities (upper and lower 20^th^ percentiles). Performance was evaluated as the difference in true error rates between these two pools of trials. **B**, Held-out prediction performance for misclassifications across joint models, single-domain test-set feature permutations (baseline shuffled, evoked shuffled), and fully shuffled empirical null baselines (100 iterations per subject). Individual participant rate differences are shown as red squares with group means overlaid. **C**, Same as **B**, for models predicting misses. **D**, Same as **B**, for models predicting false alarms.

Model condition significantly influenced prediction accuracy for all three error types (one-way repeated-measures ANOVAs, *F* > 11.4, *p* < 0.001 for all). For misclassifications, the intact model outperformed the null (one-sample *t*-test, *t*_37_ = 4.44, *p* < 0.001; **Fig. 5B**), and shuffling either baseline or evoked features significantly degraded performance (*t*_37_ = 3.75, *p* = 0.001; *t*_37_ = 2.58, *p* = 0.014, respectively). Misses, by contrast, relied on evoked dynamics alone. The intact model outperformed the null (*t*_36_ = 3.59, *p* = 0.003), shuffling the evoked feature significantly degraded prediction (*t*_36_ = 3.59, *p* = 0.003), but shuffling baseline features did not (*t*_36_ = -1.32, *p* = 0.195; **Fig. 5C**). This is consistent with our group-level findings that baseline markers did not distinguish misses from correct trials. False alarms showed a similar profile to misclassifications, with dependency on both categories of features (intact vs. null: *t*_35_ = 4.08, *p* < 0.001; baseline shuffled: *t*_35_ = 2.86, *p* = 0.007; evoked shuffled: *t*_35_ = 3.13, *p* = 0.007; **Fig. 5D**). Tightening the decision criterion for flagging high-confidence misclassification events further improved prediction margins, at the cost of increased across-subject variance (**Fig. S4**).

Additionally, we ran extended regression models which incorporated session time, run time, and inter-target intervals. Shuffling both baseline and evoked features degraded performance for all three error types, ruling out that their predictive contributions are better explained by these time-related predictors (intact extended model vs. baseline+evoked shuffled, *t* > 4.07, *p* < 0.001 for all).

Together, these predictive models demonstrate that baseline states and evoked processing failures represent complementary, non-redundant precursors to behavioral breakdown during continuous monitoring.

## DISCUSSION

Here, we investigated neural and autonomic contributions to perceptual error formation. While previous work has largely studied the underlying phenotypes of errors in isolation, typically relying on separate tasks, we developed a paradigm for studying cardinal classes of listening errors during periods of sustained listening with occasional, uncued targets. By moving away from synthetic trial-based conditions, this approach allowed us to map the pathways that underlie and shape specific error profiles before they occur.

We found that scalp-EEG negativities associated with the processing of repeating stimuli scaled systematically with upcoming behavioral outcome. This ERP signature of neural tracking mirrored the salience of the percept. For misclassification errors, the negativity was moderately decreased, indicating preserved stimulus detection but weak structural boundaries. Misses were preceded by an almost complete absence of this negativity. Conversely, false alarms were characterized by robust negativities in the neural activity that occurred in the absence of physical stimulus properties. This suggested that neural representations can mimic statistical tracking to drive phantom percepts.

Although some work suggests that certain regularity-evoked activity operates independent of attention state (Sohoglu and Chait 2016), a growing body of literature indicates it is influenced by cognitive state variables (Herrmann and Johnsrude 2018; Herrmann et al. 2019; Al Jaja et al. 2020; Herrmann et al. 2021; Ringer et al. 2023; Hu and Chait 2026). Our findings provide further evidence of high-level cognitive modulation of these responses, directly linking their variance to subsequent perceptual instability. Notably, this top-down influence did not extend to auditory steady-state responses. Although we observed strong encoding of spectral amplitude (40 Hz), we did not observe modulations in relation to subsequent outcomes, indicating stable lower-level processing. This result aligns with prior findings that neural synchronization to envelope modulation rates is largely insensitive to task-related variables (Varghese et al. 2015; Herrmann and Johnsrude 2018). Similar effects have been described in the visual system, where steady-state visual evoked potentials failed to determine upcoming lapses in vigilance that were otherwise associated with event-related activity (O’Connell et al. 2009).

Our findings suggest that multiple, at least partially separable systems contribute to error generation, spanning both baseline and evoked components of neural and physiological state. Selective shuffling of baseline and evoked features within our predictive models showed that both remained informative in forecasting upcoming errors, indicating that evoked dynamics are not simply a downstream readout of baseline state. Within the baseline window, two brain state variables, aperiodic slope and pupil-linked arousal (Donoghue et al. 2020; Lawlor et al. 2023; Parajuli et al. 2023), signaled upcoming misclassifications and false alarms, but with distinct profiles. Aperiodic slope effects appeared to differ in their topography across error types, while pupil-linked arousal showed a non-monotonic, inverted-U relationship with performance. This aligns with previous work evaluating neural synchronization and arousal states concurrently (Waschke et al. 2019; Podvalny et al. 2021), and raises the possibility that temporal tracking, cortical excitability, and arousal-related neuromodulation reflect separate, non-redundant axes of state-dependent control shaping upcoming errors (Gao et al. 2017; Grujic et al. 2024). Methodologically, this independence could also reflect individual differences in signal-to-noise ratios. Due to idiosyncrasies in anatomy and physiology, some participants may exhibit artifact-free pupillometry while others have cleaner scalp-EEG. Combining both modalities may therefore be necessary to capture these error signatures reliably across a population, rather than relying on either measure alone.

Accounting for human error in demanding, uncued listening tasks requires moving beyond static models of sensory encoding. We discovered electrophysiological and autonomic signatures of human monitoring errors that were robust across individuals. The findings hold translational relevance for identifying imminent failures in high-stakes operational environments with human operators, raising the possibility of closed-loop brain-machine interface systems designed to mitigate mistakes before they happen (deBettencourt et al. 2015; Faller et al. 2019; Meissner et al. 2023). While our cross-validated modelling highlights robust population-level signatures, there remains scope for future work to tune these models at an individual level. The physiological signatures of listening errors described here reinforce that perceptual variability is not the product of stochastic neural noise but is instead the product of empirical dynamics in underlying brain states (Resnik and Polley 2021; Ashwood et al. 2022; Hulsey et al. 2024; Papadopoulos et al. 2026). Incorporating stimulus-independent sources of brain state dynamics into models that generally consider only stimulus encoding will provide a more complete and accurate accounting of sound perception in demanding real-world listening tasks.

## ACKNOWLEDGEMENTS

This work was funded by grants from the National Institute on Deafness and Other Communication Disorders [P50DC015857 (DP)].

## Notes

### Competing Interest Statement

The authors have declared no competing interest.

## REFERENCES

Ashwood ZC et al. 2022. Mice alternate between discrete strategies during perceptual decision-making. Nature Neuroscience 2022 25:2. 25(2):201–212

Barascud N et al. 2016. Brain responses in humans reveal ideal observer-like sensitivity to complex acoustic patterns. Proceedings of the National Academy of Sciences. 113(5):E616–E625

Beerendonk L et al. 2024. A disinhibitory circuit mechanism explains a general principle of peak performance during mid-level arousal. Proceedings of the National Academy of Sciences. 121(5):e2312898121

de Cheveigné A, Simon JZ. 2008. Denoising based on spatial filtering. J Neurosci Methods. 171(2):331– 339

deBettencourt MT, et al. 2015. Closed-loop training of attention with real-time brain imaging. Nat Neurosci. 18(3):470–475

Delorme A, Makeig S. 2004. EEGLAB: an open source toolbox for analysis of single-trial EEG dynamics including independent component analysis. J Neurosci Methods. 134(1):9–21

Donoghue T et al. 2020. Parameterizing neural power spectra into periodic and aperiodic components. Nature Neuroscience 2020 23:12. 23(12):1655–1665

Eldar E, Cohen JD, Niv Y. 2013. The effects of neural gain on attention and learning. Nat Neurosci. 16(8):1146–1153

Faller J, Cummings J, Saproo S, Sajda P. 2019. Regulation of arousal via online neurofeedback improves human performance in a demanding sensory-motor task. Proc Natl Acad Sci U S A. 116(13):6482–6490

Gao R, Peterson EJ, Voytek B. 2017. Inferring synaptic excitation/inhibition balance from field potentials. Neuroimage. 158:70–78

de Gee JW et al. 2024. Strategic stabilization of arousal boosts sustained attention. Current Biology. 34(18):4114–4128.e6

Grujic N, Polania R, Burdakov D. 2024. Neurobehavioral meaning of pupil size. Neuron. 112(20):3381– 3395

Herrmann B, Araz K, Johnsrude IS. 2021. Sustained neural activity correlates with rapid perceptual learning of auditory patterns. Neuroimage. 238:118238

Herrmann B, Buckland C, Johnsrude IS. 2019. Neural signatures of temporal regularity processing in sounds differ between younger and older adults. Neurobiol Aging. 83:73–85

Herrmann B, Johnsrude IS. 2018. Neural signatures of the processing of temporal patterns in sound. Journal of Neuroscience. 38(24):5466–5477

Hu M, Chait M. 2026. Neural Signatures of Automatic Auditory Regularity Detection Reflect Individual Differences in Explicit Short-Term Memory. bioRxiv. 2026.01.09.698639

Hulsey D et al. 2024. Decision-making dynamics are predicted by arousal and uninstructed movements. Cell Rep. 43(2)

Al Jaja A, Grahn JA, Herrmann B, MacDonald PA. 2020. The effect of aging, Parkinson’s disease, and exogenous dopamine on the neural response associated with auditory regularity processing. Neurobiol Aging. 89:71–82

Lawlor J, Zagala A, Jamali S, Boubenec Y. 2023. Pupillary dynamics reflect the impact of temporal expectation on detection strategy. iScience. 106000

Maris E, Oostenveld R. 2007. Nonparametric statistical testing of EEG-and MEG-data. J Neurosci Methods. 164(1):177–190

McCormick DA, Nestvogel DB, He BJ. 2020. Neuromodulation of Brain State and Behavior. Annu Rev Neurosci. 43(Volume 43, 2020):391–415

McGinley MJ, David S V., McCormick DA. 2015. Cortical membrane potential signature of optimal states for sensory signal detection. Neuron. 87(1):179–192

Meissner SN et al. 2023. Self-regulating arousal via pupil-based biofeedback. Nat Hum Behav. 8:43–62

O’Connell RG et al. 2009. Uncovering the neural signature of lapsing attention: Electrophysiological signals predict errors up to 20 s before they occur. Journal of Neuroscience. 29(26):8604–8611

Oostenveld R, Fries P, Maris E, Schoffelen J-M. 2011. FieldTrip: Open Source Software for Advanced Analysis of MEG, EEG, and Invasive Electrophysiological Data. Comput Intell Neurosci. 2011(1):156869

Papadopoulos L et al. 2026. Modulation of metastable ensemble dynamics explains the inverted-U relationship between tone discriminability and arousal in auditory cortex. Neuron. 114(4):740–758.e19

Parajuli A, Gutnisky D, Tandon N, Dragoi V. 2023. Endogenous fluctuations in cortical state selectively enhance different modes of sensory processing in human temporal lobe. Nat Commun. 14(1):5591

Podvalny E, King LE, He BJ. 2021. Spectral signature and behavioral consequence of spontaneous shifts of pupil-linked arousal in human. Elife. 10(e68265):e68265

Resnik J, Polley DB. 2021. Cochlear neural degeneration disrupts hearing in background noise by increasing auditory cortex internal noise. Neuron. 109(6):984–996.e4

Ringer H, Schröger E, Grimm S. 2023. Perceptual learning of random acoustic patterns: Impact of temporal regularity and attention. European Journal of Neuroscience. 57(12):2112–2135

Smith SS, Sugai JA, Polley DB. 2025. Arousal state fluctuations are a source of internal noise underlying age-related declines in speech intelligibility. bioRxiv. 2025.05.09.653191

Sohoglu E, Chait M. 2016. Detecting and representing predictable structure during auditory scene analysis King AJ, editor. Elife. 5:e19113

Southwell R et al. 2017. Is predictability salient? A study of attentional capture by auditory patterns. Philosophical Transactions of the Royal Society B: Biological Sciences. 372(1714):20160105

Varghese L, Bharadwaj HM, Shinn-Cunningham BG. 2015. Evidence against attentional state modulating scalp-recorded auditory brainstem steady-state responses. Brain Res. 1626:146–164

Waschke L et al. 2021. Modality-specific tracking of attention and sensory statistics in the human electrophysiological spectral exponent. Elife. 10:e70068

Waschke L, Tune S, Obleser J. 2019. Local cortical desynchronization and pupil-linked arousal differentially shape brain states for optimal sensory performance. Elife. 8(e51501):e51501

Welch P. 1967. The use of fast Fourier transform for the estimation of power spectra: A method based on time averaging over short, modified periodograms. IEEE transactions on audio and electroacoustics. 15(2):70–73

