## SUPPLEMENTARY MATERIALS for "Neural and autonomic predictors of future listening errors"

Figures S1-S4

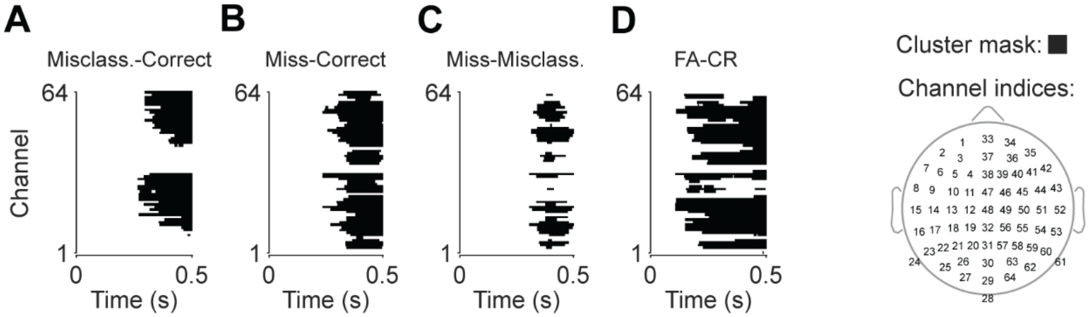


**Figure S1. ERP contrast cluster profiles.** Full spatiotemporal profiles for clusters identified in the cluster-based permutation analyses for the four ERP contrasts reported in **Fig. 2**: **A**, misclassification vs. correct; **B**, miss vs. correct; **C**, miss vs. misclassification; **D**, false alarm (FA) vs. correct rejection (CR). Black regions indicate a cluster identified as part of the cluster-based permutation analysis. Scalp topography on the right shows the corresponding channel layout.

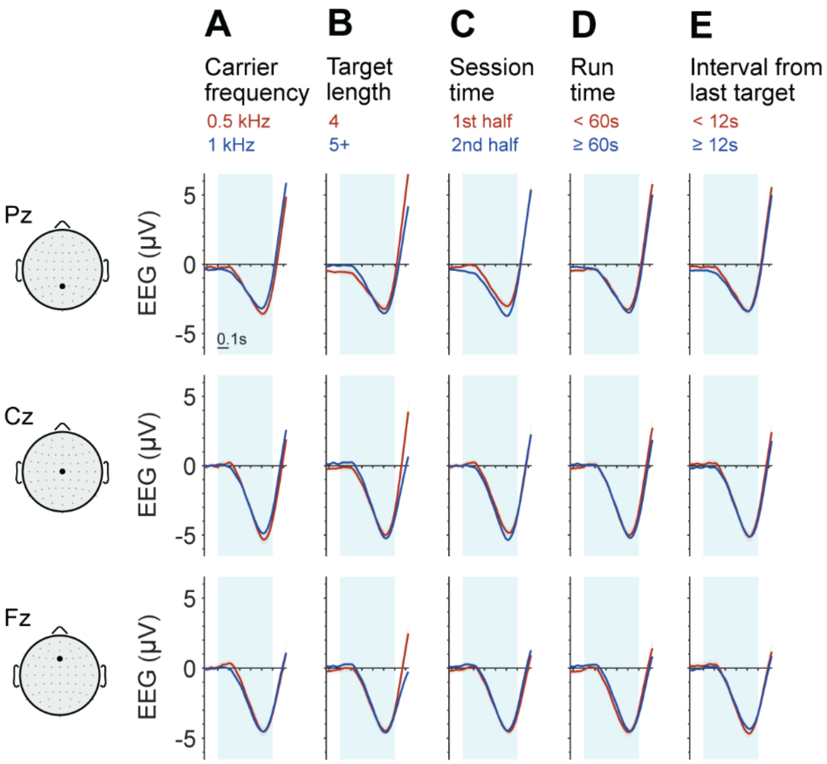


**Figure S2. No significant evoked differences for task-related contrasts.** ERPs were calculated for five different contrasts: **A,** carrier frequency, **B,** target length, **C,** session time, **D**, run time and **E**, interval since the last target. The top, middle, and bottom rows show activity at Pz, Cz and Fz electrodes, respectively. Cluster-based permutation statistical tests were performed for all contrasts, with none reaching significance.

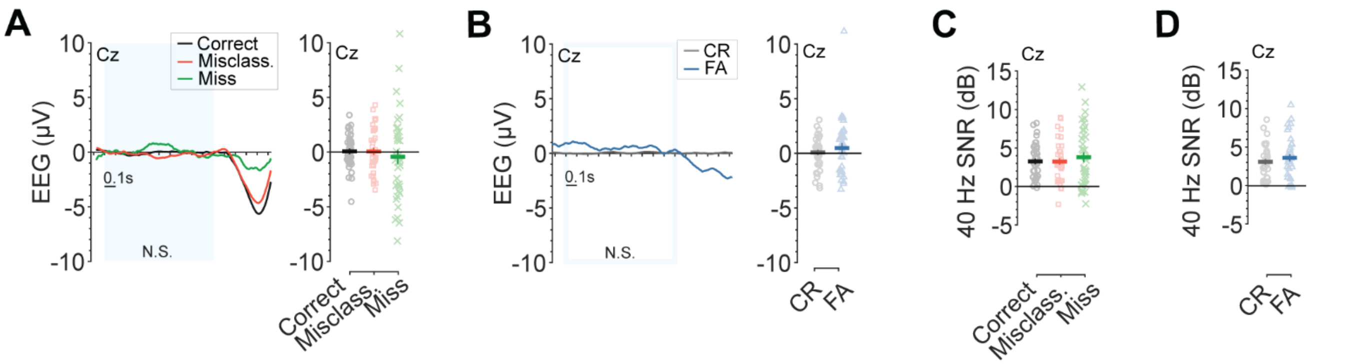


**Figure S3. ERP and ASSR analyses in the baseline window**. **A**, Grand-average ERPs at Cz in the baseline window (blue box), stratified by behavioral outcome for target presentations: correct reports (black), misclassifications (red), and misses (green). Individual-subject amplitudes are shown alongside to the right, with group means ± SEM. **B,** As in **A**, for trials without a target present: correct rejections (grey) and false alarms (blue). **C,** Individual-subject 40 Hz SNR values at electrode Cz in the baseline window, for the three target-presented outcomes: correct reports (black), misclassifications (red), and misses (green). Group means ± SEM are overlaid. **D,** As in **C**, for trials without a target present: correct rejections (grey) and false alarms (blue)

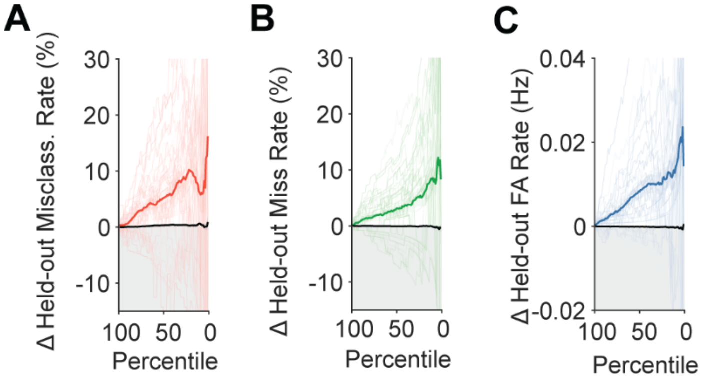


**Figure S4. Prediction margins scale with classification stringency.** Effect of decision criterion stringency on prediction margins for the intact joint model predicting **A**, misclassifications, **B**, misses, and **C**, false alarms. Stringency was varied by adjusting the percentile cutoff used to define high- and low-risk probability pools, from 1% (most stringent) to 99% (least stringent). Dark and light lines represent the sample mean and individual participants, respectively. Black lines represent the mean of the fully shuffled null distribution.
